# Unplugging lateral fenestrations of NALCN reveals a hidden drug binding site within the pore module

**DOI:** 10.1101/2023.04.12.536537

**Authors:** Katharina Schott, Samuel George Usher, Oscar Serra, Vincenzo Carnevale, Stephan Alexander Pless, Han Chow Chua

## Abstract

The sodium (Na^+^) leak channel (NALCN) is a member of the four-domain voltage-gated cation channel family that includes the prototypical voltage-gated sodium and calcium channels (Na_V_s and Ca_V_s, respectively). Unlike Na_V_s and Ca_V_s, which have four lateral fenestrations that serve as routes for lipophilic compounds to enter the central cavity to modulate channel function, NALCN has bulky residues (W311, L588, M1145 and Y1436) that block these openings. Structural data suggest that oc-cluded lateral fenestrations underlie the pharmacological resistance of NALCN to lipophilic compounds, but functional evidence is lacking. To test this hypothesis, we unplugged the fenestrations of NALCN by substituting the four aforementioned resi-dues with alanine (AAAA) and compared the effects of Na_V_, Ca_V_ and NALCN block-ers on both wild-type (WT) and AAAA channels. Most compounds behaved in a simi-lar manner on both channels, but phenytoin and 2-aminoethoxydiphenyl borate (2-APB) elicited additional, distinct responses on AAAA channels. Further experiments using single alanine mutants revealed that phenytoin and 2-APB enter the inner cav-ity through distinct fenestrations, implying structural specificity to their modes of ac-cess. Using a combination of computational and functional approaches, we identified amino acid residues critical for 2-APB activity, supporting the existence of drug bind-ing site(s) within the pore region. Intrigued by the activity of 2-APB and its ana-logues, we tested additional compounds containing the diphenylmethane/amine moiety on WT channels. We identified compounds from existing clinically used drugs that exhibited diverse activity, thus expanding the pharmacological toolbox for NALCN. While the low potencies of active compounds reiterate the resistance of NALCN to pharmacological targeting, our findings lay the foundation for rational drug design to develop NALCN modulators with refined properties.

**Significance statement:** The sodium leak channel (NALCN) is essential for survival: mutations cause life-threatening developmental disorders in humans. However, no treatment is currently available due to the resistance of NALCN to pharmacological targeting. One likely reason is that the lateral fenestrations, a common route for clinically used drugs to enter and block related ion channels, are occluded in NALCN. Using a combination of computational and functional approaches, we unplugged the fenestrations of NALCN which led us to the first molecularly defined drug binding site within the pore region. Besides that, we also identified additional NALCN modulators from existing clinically used therapeutics, thus expanding the pharmacological toolbox for this leak channel.

## Introduction

NALCN mediates a tonic Na^+^ conductance that contributes to the resting membrane potential (RMP) of excitable and non-excitable cells. Over the last two decades, ani-mal studies have shown that NALCN function regulates various bodily processes such as respiration, motor function, pain sensitivity, circadian rhythm and cancer me-tastasis (1–10). In humans, dysfunction that arises from NALCN mutations has detri-mental effects on health. Evidence from clinical studies indicates that *de novo* mis-sense mutations can cause congenital contractures of the limbs and face, resulting in characteristic facial features, hypotonia and variable degrees of developmental delay (CLIFAHDD), whereas homozygous and compound heterozygous mutations are linked to infantile hypotonia with psychomotor retardation and characteristic fa-cies (IHPRF1) (11–15). The physiological significance of NALCN has recently gar-nered interest in the ion channel field, but challenges associated with functional het-erologous expression have left NALCN lagging considerably behind other closely re-lated channels in terms of the understanding of its structure, function and pharma-cology (16–19).

As a member of the four-domain voltage-gated cation channel family, NALCN shares a common topology with the prototypical Na_V_s and Ca_V_s. These large membrane proteins are composed of four homologous but non-identical domains connected via intracellular linkers. Each of the four domains (DI–DIV) contains six transmembrane segments (S1–S6), with S1–S4 forming the voltage-sensing domains (VSDs) and S5–S6 forming the pore domains (PDs). The four VSDs are situated peripherally to a central ion-conducting pore, working in concert to couple membrane depolarisation to cation influx. Despite the conserved channel architecture, NALCN stands out from Na_V_s and Ca_V_s due to its unusual functional and pharmacological profiles. First, a prerequisite for NALCN function is the formation of a massive channelosome with three non-conducting auxiliary subunits: uncoordinated protein 79 (UNC79), uncoor-dinated protein 80 (UNC80) and family with sequence similarity 155 member A (FAM155A, also known as NALCN auxiliary factor 1 (NALF1)) (20, 21). Second, while canonical Na_V_s and Ca_V_s cycle between distinct closed, activated and inacti-vated states in response to hyperpolarisation and depolarisation of the membrane potential, NALCN shows constitutive activity that is modulated by voltage and ex-tracellular divalent cations (20). Third, NALCN exhibits strikingly low open channel probability (*P*_O_) even during periods of high activity (*P*_O_∼0.04 at −60 mV) (22). Fourth, NALCN is resistant to pharmacological targeting, with the most potent inhibitor known to date being the trivalent cation gadolinium (Gd^3+^), a promiscuous inhibitor of miscellaneous ion channels including Na_V_s, Ca_V_s, mechanosensitive, stretch-activated and transient receptor potential (TRP) channels (1, 20, 23).

In stark contrast to the lack of NALCN-specific pharmacology, Na_V_s and Ca_V_s are modulated by a vast array of natural and synthetic compounds. Animal and plant toxins generally inhibit Na_V_ and Ca_V_ function either by blocking the extracellular mouth of the ion permeation pathway or binding to the VSDs to modify channel gating (24–26). On the other hand, numerous lipophilic blockers travel across intramembrane lateral fenestrations that exist between specific interfaces of two adjacent PDs to directly occlude the central pore (**Fig. S1A**). An impressive list of clinically used drugs such as anaesthetic (e.g., lidocaine) (27), antiarrhythmic (e.g., flecainide) (28), anti-hypertensive and antianginal (e.g., verapamil and diltiazem) (29), antispasmodic (e.g., otinolium bromide) (30), and motion sickness drugs (e.g., cinnarizine) (31) enter the central cavity through these routes to inhibit Na_V_s and Ca_V_s. In addition, lateral fenestrations also house allosteric binding sites for inhibitors to negatively modulate channel function without directly blocking the pore. For example, dihydropyridine Ca_V_ blockers such as amlodipine and nifedipine bind to the fenestration between the PDs of DIII and DIV (the DIII-DIV fenestration) of Ca_V_1.1 to inhibit channel function (29, 32). Lipids are also frequent occupants of the fenestrations, with evidence suggesting that they can modulate channel function and help coordinate binding of drugs (29, 30, 33-35). Taken together, lateral fenestrations are an integral structure feature of Na_V_s and Ca_V_s that serve both as key drug access pathways as well as drug binding sites.

Since 2020, we have seen the determination of multiple structures of the NALCN channelosome (22, 23, 36-39). This nearly one-megadalton complex includes (1) the membrane-embedded NALCN, (2) the auxiliary subunit FAM155A forming a dome that sits above the channel, and (3) UNC79 and UNC80 forming a massive, intertwined superhelical assembly that docks intracellularly to the bottom of NALCN (**Fig. S1B**). The extracellular dome of FAM155A has been postulated to physically prevent molecules from accessing the selectivity filter, which may explain the insensitivity of NALCN to toxins that rely on this route to inhibit related channels (23). Despite having a central cavity with comparable volume to the classical drug-receptor site in Na_V_s and Ca_V_s, four S6 residues (W311 of DI, L588 of DII, M1145 of DIII and Y1436 of DIV) appear to plug the lateral fenestrations in NALCN (**Fig. S1C**). Based on this structural observation, an obvious question arises: could the occluded lateral fenestrations of NALCN serve as barricades to prevent drug entry, and hence contribute to the pharmacological resistance of this idiosyncratic channel? In this study, we sought to understand the impact of unplugging the lateral fenestrations on NALCN pharmacology and to expand the NALCN pharmacological toolbox.

## Results

### Alanine substitutions at the lateral gateways of NALCN widen fenestrations *in silico*

To identify the lateral fenestrations of NALCN or the lack thereof, we first subjected the WT channel structure (PDB 7SX3)(22) to an automatic tunnel detection software, MOLE 2.5 (See Materials and methods for details). While we did not identify a tunnel at the DIII-DIV interface, we found predicted lateral tunnels at the DI-DII, DII-DIII and DI-DIV interfaces of the WT channel. However, these tunnels were narrow (bottleneck radii of DI-DII=0.9 Å, DII-DIII=1.0 Å, DI-DIV=1.2 Å; **Fig. 1A**), supporting previous speculation that NALCN lacks lateral fenestrations that can serve as routes for small molecule entry into the central cavity (23). For comparison, we also analysed apo structures of Na_V_1.5 (PDB 6UZ3) and Ca_V_3.3 (PDB 7WLI). Consistent with the idea that lipids and lipophilic compounds pass through the lateral fenestrations of these channels, we detected wider lateral tunnels (bottleneck radii > 2.0 Å) at most domain interfaces (**Fig. S1D**). Next, we substituted the four key bottleneck residues of each interface of NALCN (W311 of DI, L588 of DII, M1145 of DIII, and Y1436 of DIV) with alanine *in silico* (mutagenesis performed using the PyMOL Molecular Graphics System, version 2.5.5, Schrödinger, LLC.) and applied the same analysis. This four-fold alanine (AAAA) mutant, in stark contrast to the WT channel, had lateral tunnels through DI-DII, DII-DIII, DIII-DIV and DI-DIV interfaces that were noticeably wider (**Fig. 1A**). The DI-DIV fenestration had the largest bottleneck radius of 2.7 Å, followed by DII-DIII (2.5 Å), DIII-DIV (2.2 Å) and finally DI-DII, which was the narrow-est with a bottleneck radius of 1.7 Å. To verify whether these predicted fenestration widenings are sufficient to allow compound access to the inner cavity, we proceeded to evaluate the function and pharmacology of the AAAA channel *in vitro*.

**Figure 1.**
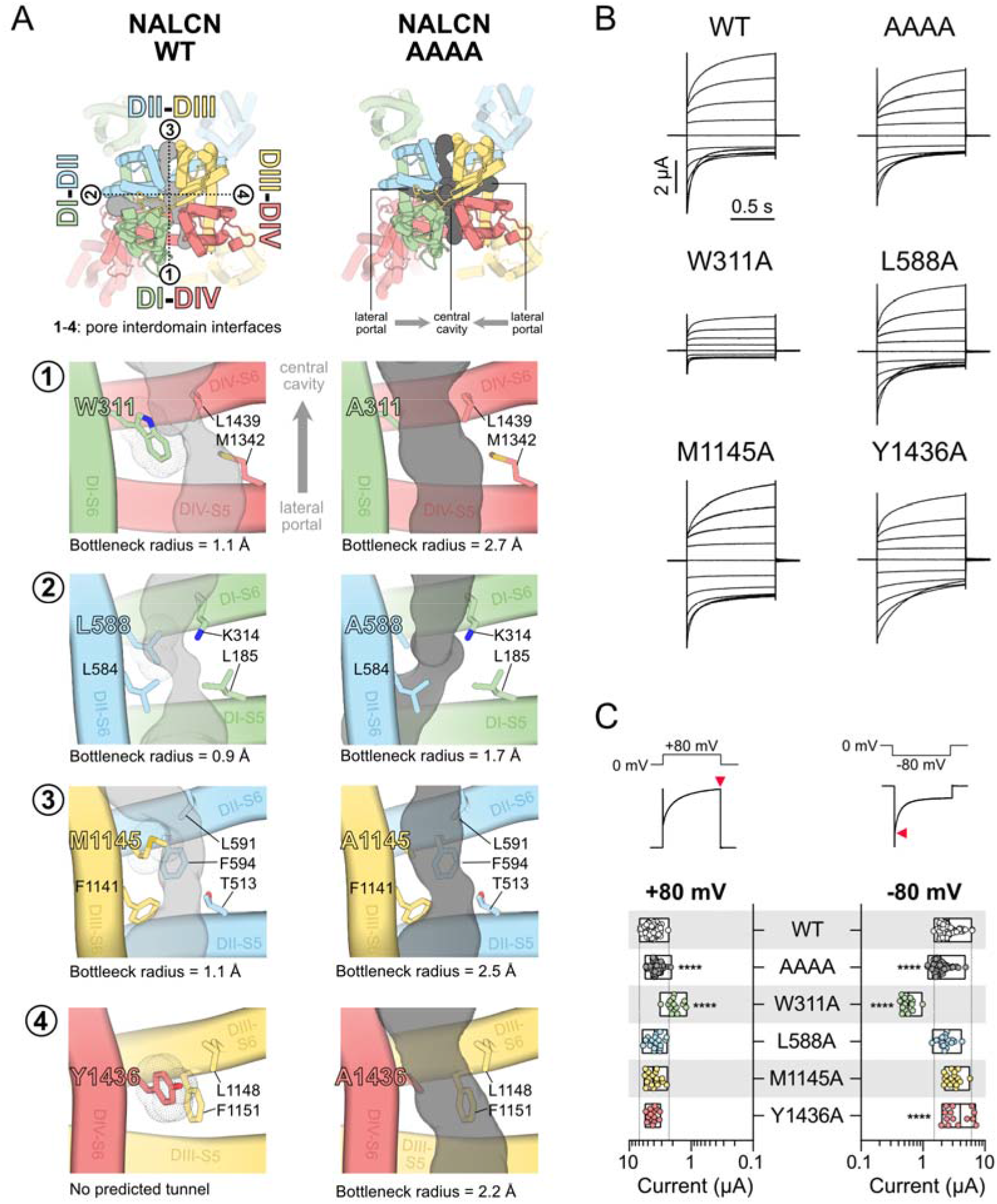
Unplugging lateral fenestrations of NALCN by substituting key bottleneck residues with alanine. (A) Top views of NALCN WT (*left*) and AAAA (*right*) channels. Predicted tunnels at I] DI-DIV, I] DI-DII, I] DII-DIII and I] DIII-DIV interfaces are shown as grey surfaces. Key bottleneck and fenestration-lining residues are labelled. The bottleneck radius values for all detected tunnels are indicated accordingly. No predicted tunnels were found at the DIII-DIV interface of WT. (B) Representative current traces from *Xenopus laevis* oocytes expressing WT, AAAA or single alanine mutant complexes (+UNC79, UNC80 and FAM155A) in response to step protocols from +80 to −100 mV (holding potential, HP=0 mV) in Ca^2+^- and Mg^2+^-free buffer. (C) The plot shows current amplitudes elicited at +80□ and −80 mV for all tested constructs (*bottom*). The current amplitudes were measured at the peak response, as indicated by red triangles (*top*). \*\*\*\**p*<□0.0001 using one-way ANOVA, Dunnett’s test (against WT). See Supplementary Table 1 for descriptive statistics. Dotted lines indicate the minimum and maximum current values of WT.

### Functional characterisation of the alanine mutants in *Xenopus laevis* oocytes

To determine the effect of these alanine mutations on channel function, we introduced single (W311A, L588A, M1145A and Y1436A) and four-fold alanine mutations into NALCN using site-directed mutagenesis and expressed these mutant channels with the auxiliary subunits UNC79, UNC80 and FAM155A in *Xenopus laevis* oocytes. We then measured voltage-evoked currents from each construct by two-electrode voltage-clamp five days post RNA injection. The WT NALCN channel conducted robust currents in response to depolarisation and hyperpolarisation from a holding potential of 0 mV (4.4±0.9 µA at +80 mV; 2.6±1.0 µA at −80 mV; *n*=73; **Fig. 1B** and **C**). The AAAA mutant channel showed a small but significant decrease in current amplitudes (3.7±0.6 µA at +80 mV; 1.9±0.6 µA at −80 mV; *n*=75; *p*<0.0001; one-way ANOVA, Dunnett’s test (compared to WT); **Suppl. Table 1**; **Fig. 1B** and **C**). The single W311A mutation resulted in markedly lower current amplitudes both in the outward and inward directions (1.9±0.5 µA at +80 mV; 0.6±0.1 µA at −80 mV; *n*=14; *p*<0.0001; one-way ANOVA, Dunnett’s test (compared to WT); **Suppl. Table 1**; **Fig. 1B** and **C**). The remaining three single mutants L588A, M1145A and Y1436A behaved similarly to WT channels, with no significant differences in current amplitudes, except for a slight increase for Y1436A at hyperpolarised potentials (4.0±1.9 µA at −80 mV; *n*=14; *p*<0.0001; one-way ANOVA, Dunnett’s test (compared to WT); **Suppl. Table 1**; **Fig. 1B** and **C**). Consistent with observations from disease-causing mutations in the NALCN pore (23), we also observed significant differences in the current deactivation kinetics in response to hyperpolarizing voltage steps with the L588A, M1145A and Y1436A mutants (**Suppl. Table 1**; **Fig. S2**). However, we did not investigate this effect further.

### Pharmacological screening reveals distinct responses for phenytoin and 2-APB on WT and AAAA channels

Having established that the AAAA mutant channel displayed WT-like function, we evaluated if the fenestrations or the central cavity of NALCN houses potential drug binding sites. We hypothesised that by widening the lateral fenestrations of NALCN, some compounds that are ineffective on WT channel would exert modulatory effect on the AAAA channel. Alternatively, previously reported inhibitors of NALCN may have increased potency or additional, previously unseen effects on the mutant channel. To test this hypothesis, we expressed both WT and AAAA channel complexes in *Xenopus laevis* oocytes and then measured their responses to 13 lipophilic channel modulators. We chose these compounds either for their ability to enter Na_V_s or Ca_V_s via the lateral fenestrations or for their reported inhibitory effect on NALCN. Considering the pharmacological resistance of NALCN to previously tested compounds, we started testing each compound with the maximum soluble concentration achievable or 1 mM if solubility was not an issue (concentration range=100 µM to 1 mM).

Overall, the effects of these compounds on both WT and AAAA channels were diverse (**Fig. 2A** and **B**). Five out of the 13 compounds tested (carbamazepine, lidocaine, phenytoin, lacosamide, and Z944) showed no effect on WT channels; two selectively inhibited the outward (CP96345) or inward (lamotrigine) currents; three inhibited the outward and potentiated the inward currents (quinidine, diltiazem, and propafenone); three inhibited both outward and inward currents (nifedipine, L-703,606, and 2-APB). In addition to the effects on current amplitudes, we also observed changes in current phenotype. For example, the application of quinidine, diltiazem, propafenone and CP96435 resulted in a “hooked” current at hyperpolarized potentials, suggesting that these compounds may have an impact on channel gating (**Fig. 2A**).

**Figure 2.**
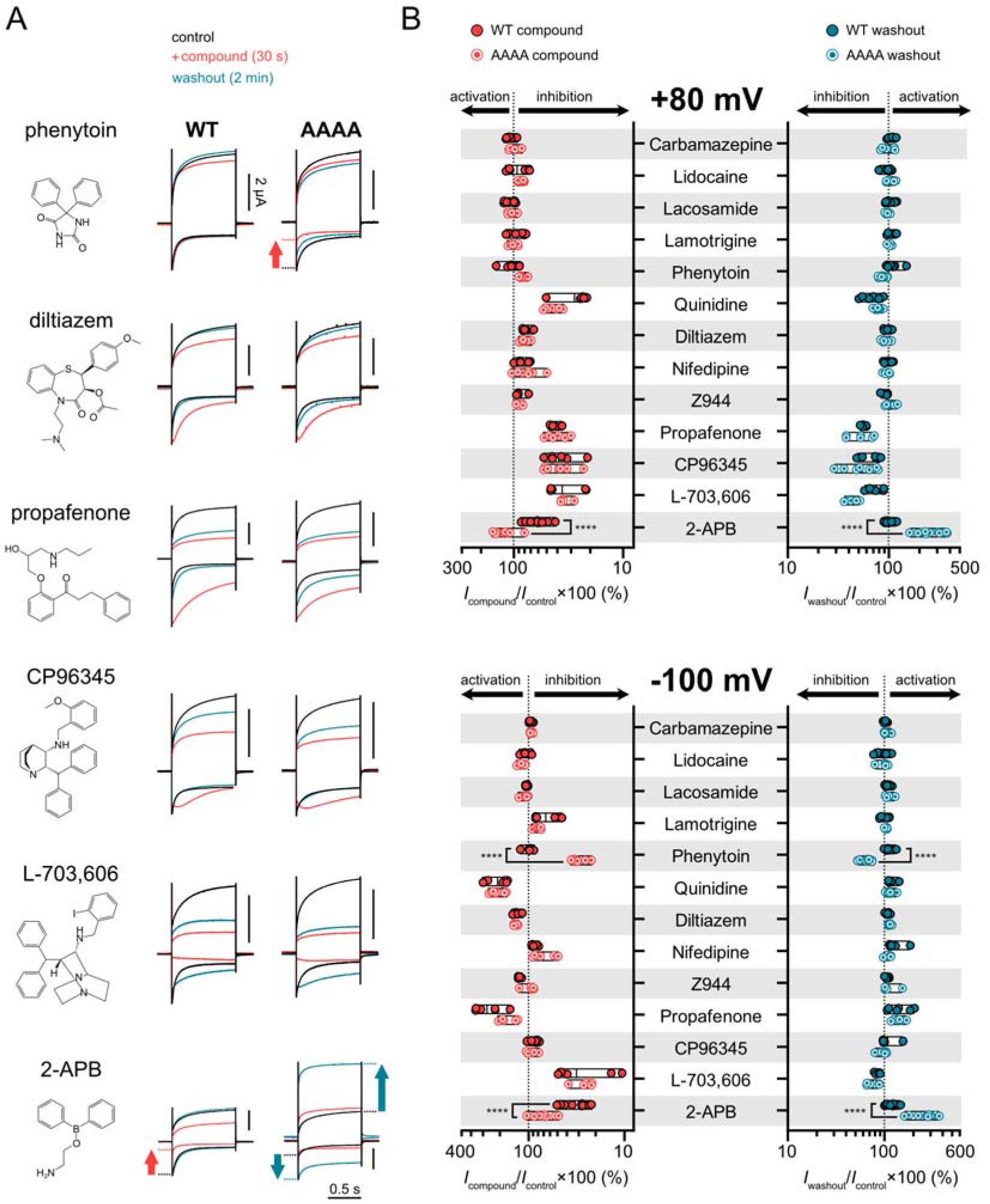
Effects of Na_V_, Ca_V_ and NALCN inhibitors at NALCN WT and AAAA channels. (A) Representative current traces from *Xenopus laevis* oocytes express-ing WT or AAAA channel complexes (+UNC79, UNC80 and FAM155A) during con-trol (black), compound application (red) and washout (blue) at +80 (outward) and −100 mV (inward) from a HP of 0 mV. The inhibitory effects of phenytoin and 2-APB at −100 mV are highlighted with red arrows, whereas the post-washout activation effects of 2-APB at +80 and −100 mV are highlighted with blue arrows. (B) The plots show efficacy of different compounds at WT and AAAA channels during application (red) and washout (blue) normalised against control current at +80 mV (top) and −100 mV (bottom). \*\*\*\**p*<□0.0001 using unpaired t-test.

The AAAA mutant channel responded similarly to the WT channel in response to most of these compounds, except for phenytoin and 2-APB. While the application of 300 µM phenytoin had no effect on WT channel currents, it significantly inhibited the inward current of AAAA channels (64.8±6.7 % inhibition at −100 mV; *n*=13; *p*<0.0001; unpaired t test; **Fig. 2A** and **B**). We have previously reported the inhibitory effect of 2-APB on NALCN (23). On WT channels, 1 mM 2-APB had a more prominent effect on inward (70.2±8.2 % inhibition at −100 mV; *n*=17) than outward current (23.2±26.7 % inhibition at +80 mV). The blocking effect of 2-APB was reversible, evident from the full recovery of currents after washout for two minutes (**Fig. 2A**). By contrast, 1 mM 2-APB only weakly inhibited the inward current of the AAAA mutant (32.9±12.9 % inhibition at −100 mV; *n*=21; **Fig. 2B**). We observed slight activation of the outward current (30.5±27.0 % of activation at +80 mV). To our surprise, following the removal of 2-APB with a two-minute washout, the AAAA mutant channel showed even greater activation in both inward (167.9±50.3 %) and outward (227.7±122.4 %) direc-tions.

### Phenytoin inhibits M1145A and 2-APB activates W311A

To determine if phenytoin and 2-APB elicited their unique responses at AAAA channels via specific lateral fenestrations, we tested these compounds on individual alanine mutants. Like WT channels, the W311A, L588A and Y1436A mutants did not react to 300 µM phenytoin (**Fig. 3A**). The M1145A mutant, on the other hand, showed noticeably reduced currents in response to phenytoin (31.0±9.2 % of inhibition at −100 mV; *n*=11; *p*<0.0001; one-way ANOVA, Dunnett’s test (compared to WT); **Fig. 3A**). For 2-APB, the reversible inhibitory effect observed on WT channels (**Fig. 2**) was replicated with L588A, M1145A and Y1436A mutants (**Fig. 3B**). By contrast, 2-APB activated W311A readily during application (87.1±60.2 % and 143.3±92.5 % of activation at +80 and −100 mV, respectively; *n*=9; **Fig. 3B**). This stimulatory effect was even more pronounced post washout, with 334.6±146.6 % and 484.0±238.7 % of activation measured at +80 and −100 mV, respectively. Taken together, these results suggest that these compounds access their putative binding site(s) via different lateral fenestrations, with phenytoin and 2-APB likely using the DII-DIII and DI-DIV fenestrations, respectively.

**Figure 3.**
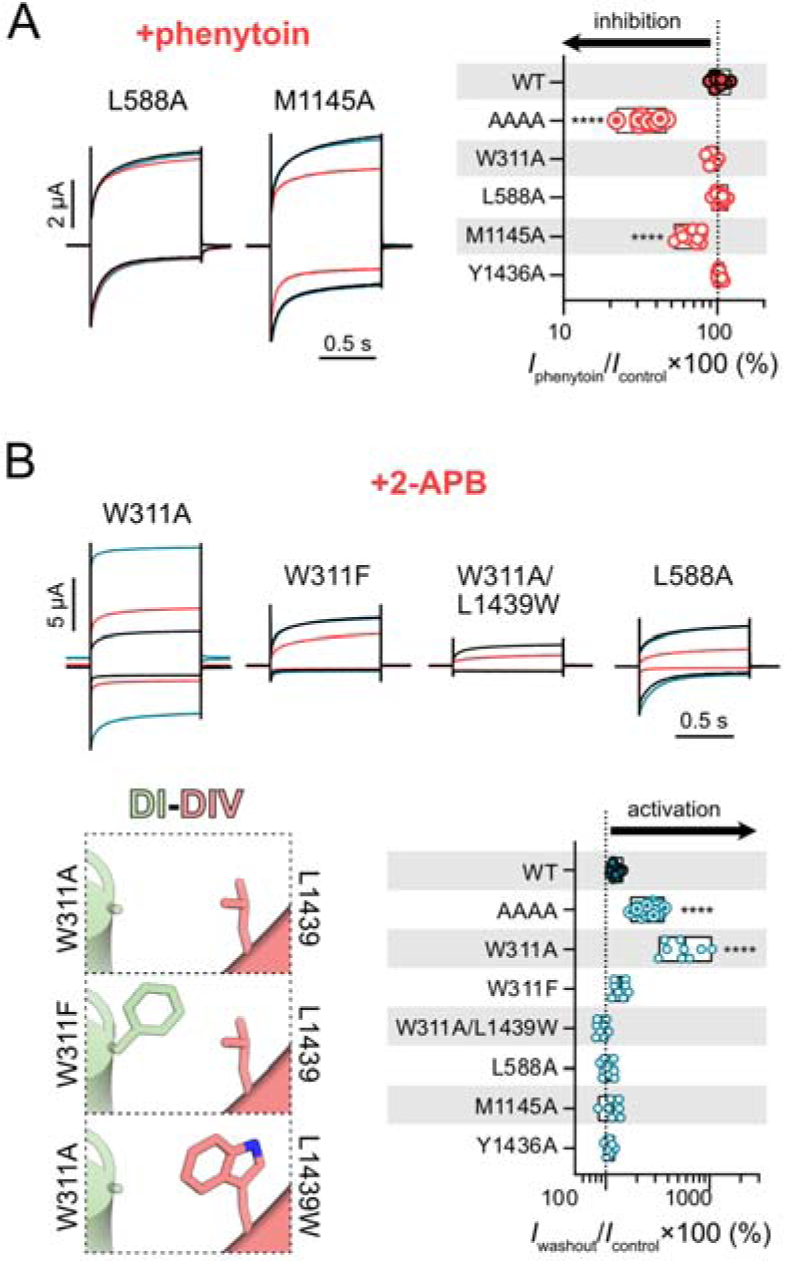
Phenytoin and 2-APB modulate NALCN AAAA via distinct unplugged fenes trations. (A) *Left*, representative current traces showing the inhibitory effect of ¼M pheny toin on the L588A and M1145A mutant chan nels at +80 (outward) and −100 mV (inward) from a HP of 0 mV. *Right*, the plot shows per centage of current left for NALCN WT and dif ferent mutants during the application of pheny toin normalised against control current elicited at +100 mV. (B) *Top*, representative current traces from *Xenopus laevis* oocytes expressing different mutants in response to application (red) and washout (blue) of 1 mM 2-APB in re sponse to step protocols at +80 (outward) and −100 mV (inward) from a HP of 0 mV. *Bottom left*, side view of the DI-DIV fenestrations of W311A, W311F and W311A/L1439W mutants, highlighting the side chains of position (DI) and 1439 (DIV). *Bottom right*, the plot shows percentage of current left for NALCN WT and different mutants post 2-APB washout normaised against control current elicited at −100 mV. \*\*\*\**p*<□0.0001 using one-way ANOVA, Dun nett’s test (against WT). See Supplementary Table 1 for descriptive statistics.

The W311A mutation was predicted to widen the DI-DIV fenestration (**Fig. 1A**). If a wider DI-DIV fenestration was directly responsible for the 2-APB activation observed at W311A and AAAA, we hypothesised that reintroducing a bulky residue at the DI-DIV interface of the W311A mutant should prevent 2-APB entry and activation. For this purpose, we generated two new mutants W311F and W311A/L1439W. These mutants had narrower predicted tunnels at the DI-DIV interface (bottleneck radii of 1.3 and 1.6 Å for W311F and W311A/L1439W, respectively; **Fig. 3B**). Like W311A, both mutants showed reduced current amplitudes compared to WT, suggesting that tryptophan at position 311 plays a critical role for channel function as even a subtle tryptophan- to-phenylalanine substitution was not well tolerated. The application and subsequent removal of 2-APB, however, did not result in current activation at these mutants (**Fig. 3B**), supporting our speculation that 2-APB is able to enter and bind to a previously inaccessible activation site only when we unplugged the DI-DIV fenes-tration with the W311A mutation.

### Inhibition masks activation during 2-APB application

We were intrigued by the delayed activation effect of 2-APB at W311A and AAAA channels and wanted to further investigate the reversibility (permanent or washable) and reason behind the post-washout prominence of this effect. To this end, we repeated the 2-APB experiments on the AAAA mutant but measured currents in two-minute intervals after the first washout to determine the time required for currents to return to baseline. We found that, despite the removal of 2-APB, it took as long as eight minutes for the current responses to gradually return to baseline (**Fig. 4A**, left panel). This slow washout profile is consistent with the existence of an activation site for 2-APB buried deep within the pore of W311A and AAAA channels. By contrast, the quick washout of the inhibitory effect of 2-APB on WT (**Fig. 2A**) is consistent with an extracellular, solvent-accessible inhibitory binding site. This two-site model can account for the delayed appearance of activation as co-occupation of both the inhibition and activation sites can mask activation during 2-APB application (**Fig. 4B**). To test this hypothesis, we reapplied 2-APB for 30 seconds after the two-minute wash-out period and observed immediate return of the elevated current to baseline levels (**Fig. 4A**, right panel), indicating that inhibition and activation occur simultaneously during 2-APB perfusion. Considering the anticipated rapid departure of 2-APB from the solvent-accessible inhibitory site upon washout, the heightened current response observed during this step can be explained by the sustained binding or slow unbinding of 2-APB at the activation site within the pore. abolished the activation effect at −100 mV, whereas Q279A and S592A mutations had no discernible effects on 2-APB’s actions on NALCN AAAA. The inhibitory effect of 2-APB was still observed for the alanine mutants that do not show post-washout activation, corroborating our hypothesis that inhibition and activation are mediated via two distinct binding sites.

**Figure 4.**
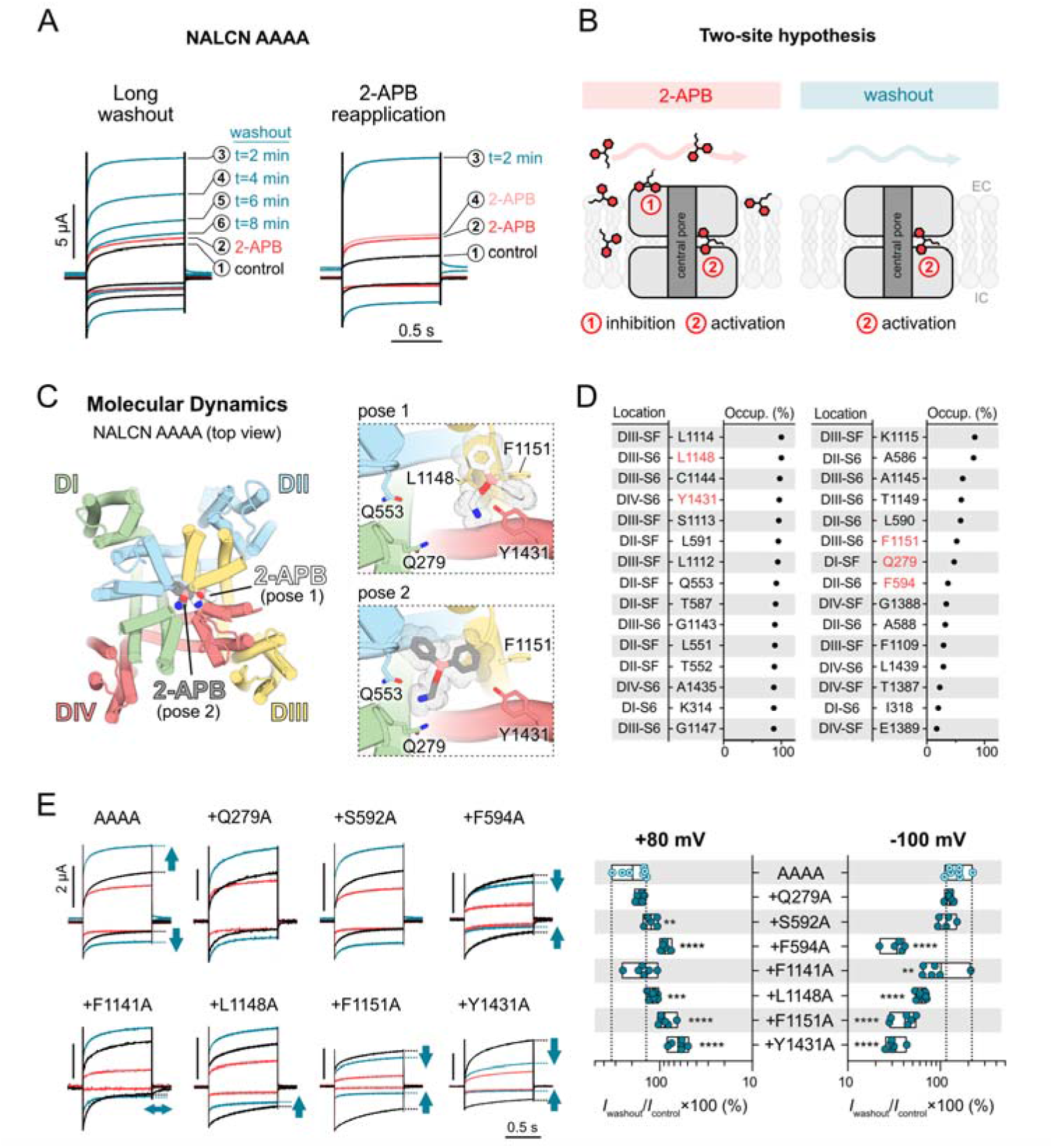
Locating the putative activation site of 2-APB in the NALCN pore. (A) Representative current traces from *Xenopus laevis* oocytes expressing the AAAA mutant in response to application of 1 mM 2-APB using two different protocols: *left*, single application of 2-APB (□) and washout for 8 min with currents measured at 2-min intervals (□–□); *right*, applications of 2-APB before (□) and after (□) a single 2-min washout step (□). (B) Schematic of our proposed two-site model for 2-APB at NALCN AAAA channels. (C) *Left*, molecular dynamics simulations using NALCN AAAA led to the two most common binding poses of 2-APB (white and grey). *Right*, zoomed views of binding pose 1 (white) and 2 (grey) of 2-APB, highlighting key resi-dues in the vicinity of 2-APB. (D) Top 30 residues showing the highest percentage occupancy (occup.), *i.e.*, the percentage of time a residue was within a 5 Å proximity to 2-APB during the ∼50-ns trajectory. The location of each residue is provided (SF: selectivity filter), and the residues highlighted in red were chosen for mutagenesis and functional evaluation. (E) *Left*, representative current traces from *Xenopus laevis* oocytes expressing various mutant channels (on the background of NALCN AAAA) in response to application of 1 mM 2-APB. *Right*, the plot shows percentage of cur-rent left for NALCN AAAA and different mutants post 2-APB washout normalised against control current elicited at −100 mV. \*\**p*<0.01; \*\*\**p*<0.001; \*\*\*\**p*<□0.0001 us-ing one-way ANOVA, Dunnett’s test (against AAAA). See Supplementary Table 1 for descriptive statistics.

### Alanine substitutions of hydrophobic residues in the inner cavity abolish 2-APB activation

To test the hypothesis that there is a binding site for 2-APB in the inner cavity of NALCN AAAA, we turned to computational approaches. First, we performed unbi-ased MD simulations to elucidate the dynamic binding patterns of 2-APB within the inner cavity of NALCN AAAA (**Fig. 4C**). Two common binding configurations of 2-APB emerged from these simulations, one involving multiple residues located around the DIII-DIV fenestration (pose 1) and another at the bottom of the selectivity filter (pose 2; **Fig. 4C**). In parallel, we performed molecular docking using 2-APB and phenytoin within a defined search space encompassing the inner cavity and lateral fenestrations of NALCN AAAA (**Fig. S3A**). We found that majority of docked poses of 2-APB align with pose 1 of unbiased MD simulations, near a binding pocket formed by L1148, F1151 (DIII) and Y1431 (DIV). By contrast, phenytoin appears to be binding in the middle of the ion permeation pathway, which may explain its inhibi-tory action on NALCN AAAA (**Fig. S3A**).

Over the course of these simulations, we noticed that some of the 2-APB-interacting residues suggested by MD simulations and molecular docking are homologous to the binding residues of verapamil and cinnarizine at Ca_V_1.1 and 1.3, respectively (**Fig. S3B**). Guided by our computational predictions and the well-established drug binding sites in the pores of structurally related Ca_V_ channels, we chose seven po-tential 2-APB-contacting residues (Q279, S592, F594, F1141, L1148, F1151 and Y1431) and substituted these residues with alanine on the NALCN AAAA back-ground. All mutants were functional upon injection into *Xenopus* oocytes. We then measured current responses at +80 and −100 mV during the application and washout of 2-APB (**Fig. 4E**). Three mutations (F594A, F1151A and Y1431A) completely abol-ished the post-washout activation effect of 2-APB on NALCN AAAA, two mutations (F1141A and L1148A) only

### Single extracellular mutations do not affect 2-APB inhibition

To test the hypothesis that the inhibitory binding site(s) are on the extracellular surface of the channel, we first chose several potential interacting residues based on the cryo-EM structure of 2-APB-bound TRPV3 channel (**Fig. S4A**), where hydrophobic and hydrophilic residues (Y540, R487 and Q483) interact with 2-APB, and mutations of these residues affect the potency of 2-APB action at TRPV3 (40). We thus substituted eight hydrophobic and charged residues on the extracellular side of the four non-identical NALCN VSDs (VSD1: F61, E128; VSD2: R412, Y470; VSD3: R911; VSD4: H1291, Y1300) and the pore domain (Y1412) with alanine (**Fig. S4A**). On the background of NALCN AAAA, we anticipated these mutations to affect inhibition and allow prominent activation to be observed during 2-APB application (instead of after washout). All eight mutants were functional upon injection. However, all mutants behaved similarly to AAAA (**Fig. S4B**), suggesting that these single alanine mutations did not affect 2-APB inhibition. Therefore, the inhibitory 2-APB site remains elusive.

### Effects of different 2-APB analogues on NALCN WT channels

Owing to the ability of 2-APB to form a nitrogen-to-boron coordinate covalent bond, the molecule can exist in different forms including a ring form, an open chain form or a dimeric form (**Fig. 5A**) (41). Our data already showed that phenytoin, a structural analogue of the ring form, did not have any effect on NALCN WT currents even when applied at concentration as high as 300 µM (**Fig. 2** and **5B**). Considering the ability of 2-APB to switch between its different forms, we investigated which form(s) of 2-APB were responsible for the inhibition of WT currents. We therefore tested the effects of two 2-APB analogues on WT channels. The antihistamine diphenhydramine resembles the open chain form of 2-APB (**Fig. 5A**). At 1 mM, diphenhydramine inhibited the outward current (60.3±4.3 % of inhibition at +80 mV; *n*=9; **Fig. 5B**) and activated the inward current (84.9±19.8 % of activation at −100 mV; *n*=9; **Fig. 5B**). The dimeric analogue diphenylboronic anhydride (DPBA), on the other hand, inhibited NALCN current in both directions with high efficacy at 1 mM (66.9±4.0 % of inhibition at +80 mV; *n*=7; *p*<0.0001; one-way ANOVA, Dunnett’s test (compared to 2-APB); **Suppl. Table 1**; **Fig. 5B**). All in all, our results indicate that the distinct forms of 2-APB affect NALCN function differently: the dimeric form of 2-APB (represented by DPBA) is the most efficacious in inhibiting NALCN function, the monomeric ring form (represented by phenytoin) has no detectable effect on NALCN function, whereas the open-chain monomeric form (represented by diphenhydramine) shows mixed inhibitor/activator effects in a voltage-dependent manner.

**Figure 5.**
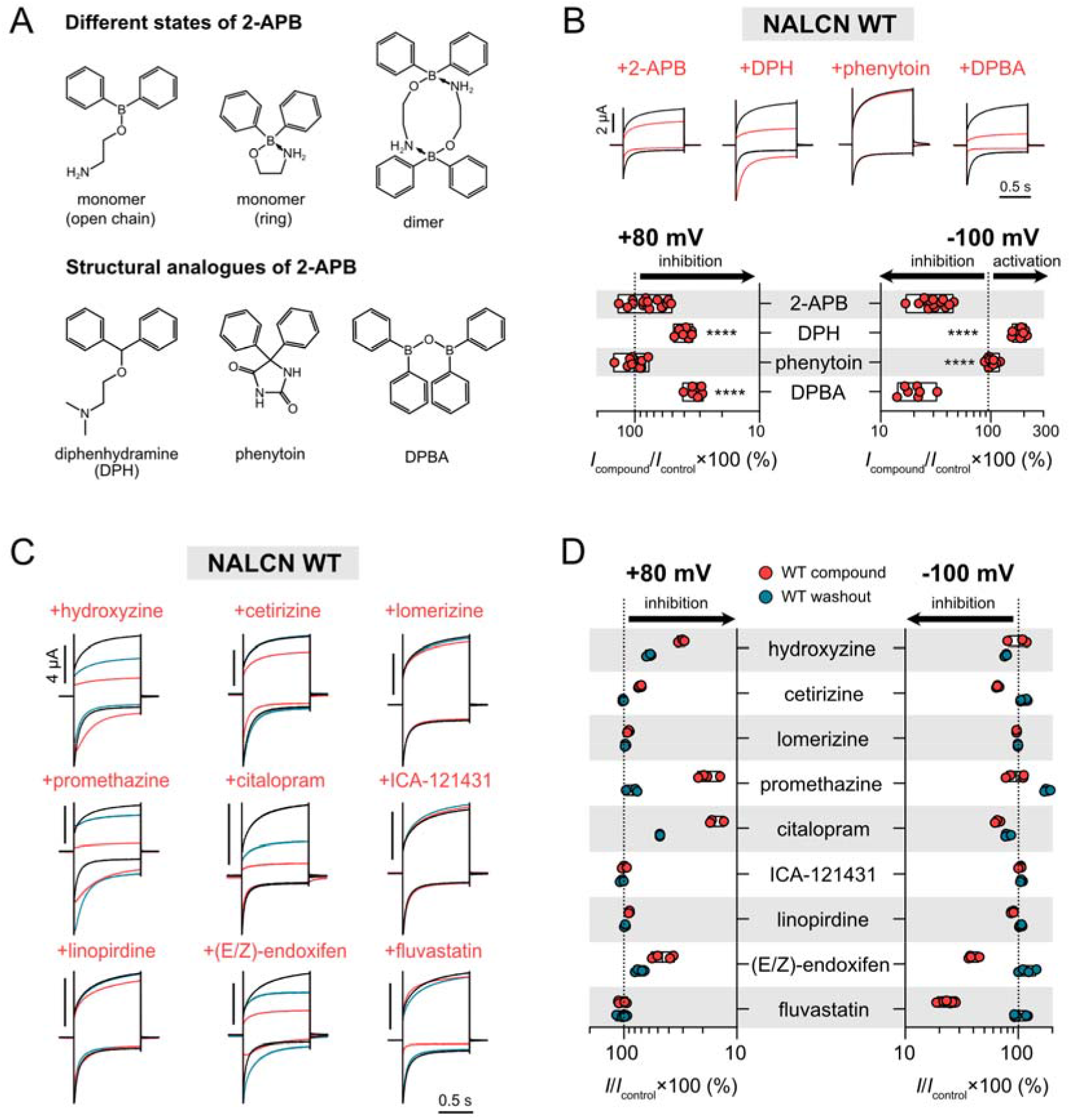
Effects of 2-APB analogues and compounds containing the di-phenylmethane/amine motif on NALCN WT channels. (A) Different forms of 2-APB (*top*) and their corresponding structural analogues (*bottom*). (B) *Top,* represen-tative current traces from *Xenopus laevis* oocytes expressing NALCN WT channel complex (+UNC79, UNC80 and FAM155A) in response to 1 mM 2-APB, 1 mM di-phenhydramine (DPH), 300 µM phenytoin or 1 mM diphenylboronic anhydride (DPBA) at +80 mV (outward) and −100 mV (inward). *Bottom*, the plot shows percent-age of current left during the application of 2-APB, DPH, phenytoin and DPBA. \*\*\*\**p*<□0.0001 using one-way ANOVA, Dunnett’s test (against 2-APB). See Supple-mentary Table 1 for descriptive statistics. (C) Representative current traces from *Xenopus laevis* oocytes expressing NALCN WT channel complex in response to dif-ferent compounds (see Methods for concentrations tested). (D) The plots show effi-cacy of different compounds at WT NALCN channels during application (red) and washout (blue) normalised against control current at +80 mV (top) and −100 mV (bot-tom).

### Functional screening of compounds containing the diphenylmethane/amine motif

Up until this point, we found only three small molecules that have strong inhibitory effect on WT channel, namely L-703,606, 2-APB and DPBA (∼70–80 % of inward current inhibition at −100 mV; **Fig. 2B** and **5B**). However, none of these compounds had potency comparable to that of the trivalent cation Gd^3+^. Since a common feature of these compounds is the diphenyl-x (x=carbon/boron) motif, we tested if other small molecule drugs with similar feature also modulate NALCN function. For this purpose, we tested nine compounds that have the diphenylmethane/amine motif (**Fig. S5**) including antihistamines (hydroxyzine, cetirizine, lomerizine and promethazine), antidepressants (citalopram), cognitive-enhancing drugs (linopirdine), anticancers ((E/Z)-endoxifen), statins (fluvastatin) and pharmacological tools (ICA−121431, a potent blocker of Na_V_1.1 and Na_V_1.3) on WT channels. We found that three compounds (lomerizine, ICA−121431 and linopirdine) had little to no effect on NALCN when applied at 100 µM (**Fig. 5C** and **D**), three (hydroxyzine, promethazine and citalopram) had a stronger inhibitory effect on the outward than inward current, two (cetirizine and (E/Z)-endoxifen) inhibited both outward and inward currents, and one (fluvastatin) had no effect on the outward current, but inhibited the inward current efficaciously. These data suggest that the diphenyl-x (x=carbon/boron/nitrogen) motif may be a promising starting point for future structure-activity relationship stud-ies.

## Discussion

In 1977, in an attempt to explain the ability of neutral local anaesthetics to inhibit Na_V_s even when the intracellular gate is closed, Hille proposed that there are alternative hydrophobic pathways in the membrane for lipid-soluble blockers to “come and go from the receptor” (42). The existence of such pathways has since been overwhelmingly supported by functional, structural and computational studies (27, 30, 34, 43-47). In eukaryotic Na_V_s and Ca_V_s, four lateral fenestrations that extend from the cell membrane to the inner pore exist at the interfaces between the S5 and S6 helices of neighbouring PDs (DI-DIV, DI-DII, DII-DIII and DIII-DIV; **Fig. S1A**) (48). These fenestrations are hydrophobic in nature and the physical dimensions of each fenestration are unique and dynamic, as they change depending on the functional state of the channel and the presence of lipids or inhibitors (46, 49-51). Functionally, these side passages serve both as routes for lipids and lipophilic molecules to enter or leave the central cavity and as allosteric binding sites. Additionally, they are potential sites for drug-drug interactions, as exemplified by the concomitant use of the antiarrhythmic agent amiodarone and the antiviral sofosbuvir (52). Amiodarone inhib-its Ca_V_ function by binding to the DIII-DIV fenestrations of the Ca_V_1 subfamily mem-bers, but its binding unexpectedly helps anchor the binding of sofosbuvir in the central cavity, leading to synergistic pore block and fatal heartbeat slowing (52). There is also increasing appreciation for the pathophysiological relevance of lateral fenestrations following reports of disease mutations identified in this region of Na_V_s (48). These mutations not only affect intrinsic channel function, but also alter the physical and/or chemical nature of these fenestrations, which may in turn affect drug accessibility via these routes. Hence, a detailed understanding of the properties and drug-gability of these often-overlooked side passages will help elucidate their roles in diseases and their potential exploitation in clinical or medical settings.

### Lateral fenestrations are key functional regions of NALCN

In this study, we show that alanine substitutions of four bulky residues (W311, L588, M1145 and Y1436) that plug the lateral fenestrations of NALCN affect channel function to varying degree: W311A reduces current amplitudes considerably (**Fig. 1C**), whereas L588A, M1145A and Y1436A mutations significantly slow current deactivation kinetics at hyperpolarizing potentials, with M1145A and Y1436A having more pronounced effects (**Fig. S2**). W311 appears to be critical for channel function as even subtle changes to the physicochemical properties of this residue, e.g., when substituted with phenylalanine, are detrimental for channel function (W311F; **Fig. 3B**). While we have not assessed the impact of these mutations on protein expression levels using biochemical methods, 2-APB’s ability to activate W311A and AAAA channels to current levels comparable to those of WT channels (**Fig. 2A** and **3B**) suggest a gating defect rather than trafficking and/or expression issues. W311, L588, M1145 and Y1436 have previously been shown to stabilise the π-bulges of the S6 helices (23), a structural feature that is gaining recognition as the gating hinge of structurally related ion channels (40, 53). As NALCN gating is still poorly understood, we speculate that alanine mutations of these residues may destabilise these critical hinge regions, impairing individual S6 movements in a domain-dependent manner, leading to either pore collapse (manifested as a dramatic reduction in currents as seen in the case of W311A) or delayed closure of the activation gate during repolarization (manifested as slower deactivation kinetics as observed in the L588A, M1145A and Y1436A mutations). All in all, our data support the emerging concept that lateral fenestrations are not just simple pathways for drug and lipid binding but are also key functional regions that contribute to channel gating.

### Potential drug binding sites in and beyond the plugged lateral fenestrations of NALCN

Thus far, all six available structures of NALCN lack accessible lateral fenestrations, which is evident from the narrow bottleneck radii and the absence of resolved lipid molecule densities in this region (22, 23, 36-39). Lateral fenestrations contribute immensely to the pharmacology of the four-domain voltage-gated cation channel family with various clinical and pharmacological compounds detected in specific individual fenestrations of Na_V_ and Ca_V_ cryo-EM structures (e.g., bulleyaconitine A: DI-DII of Na_V_1.3; flecainide: DII-DIII of Na_V_1.5; nifedipine: DIII-DIV of Ca_V_1.1; A−803467: DI-DIV of Na_V_1.8; **Fig. S1A**) (28, 29, 54, 55). We hypothesised that the lack of accessible lateral fenestrations may contribute to the pharmacological resistance of NALCN. Here, we demonstrate that substituting key bulky bottleneck residues that block these fenestrations in NALCN with alanine, which possesses a smaller side chain, renders the mutant channels sensitive to pharmacological effects not observed with WT channels. A particularly striking example is phenytoin, which clearly inhibits inward currents of AAAA channels, but has negligible activity on WT channels (**Fig. 2A** and **B**). On the other hand, 2-APB inhibits both WT and AAAA channels, but is also able to activate AAAA channels simultaneously.

To determine if phenytoin and 2-APB have preferred entry route(s) at the AAAA mutant, we tested the effects of both compounds at single alanine mutant channels that have only one fenestration unplugged at a time. Our results show that phenytoin selectively inhibits the single M1145A mutant (**Fig. 3A**), whereas 2-APB selectively activates the single W311A mutant (**Fig. 3B**). These data are consistent with our hypothesis: plugged lateral fenestrations contribute to the pharmacological resistance of NALCN, and freeing up these blocked routes allows drug entry, resulting in new pharmacological effects. In addition, they suggest that phenytoin and 2-APB favour the DII-DIII and DI-DIV fenestrations, respectively, which are the widest fenestrations based on our *in silico* predictions (**Fig. 1A**).

### The existence of 2-APB binding site(s) in AAAA channels

The synthetic compound 2-APB is a membrane-permeable, multi-target modulator that inhibits inositol 1,4,5-trisphosphate (IP3) receptors (56), activates two-pore potassium (K2P) channels (57), and inhibits/activates Ca^2+^ release-activated Ca^2+^ (CRAC) channels (58) and different members of the Transient Receptor Potential (TRP) channel family (59). We have previously reported 2-APB as a low-potency in-hibitor of NALCN (23). Here, we also find that 2-APB activates NALCN efficaciously when the DI-DIV fenestration is unplugged (**Fig. 2A** and **3B**). To explain our functional data, we propose a two-site model for 2-APB in which the inhibitory binding site(s) are found on the extracellular side of NALCN, whereas the activation site(s) are found in the inner cavity (**Fig. 4B**). During 2-APB application, the compound binds simultaneously to the inhibitory and activation sites at the AAAA or W311A mutant, masking the activation effect of 2-APB during drug application. Following drug removal with a two-minute buffer washout, 2-APB leaves the extracellular inhibitory site rapidly. However, 2-APB stays bound at the activation site due to the lipophilicity of both compound and fenestration, resulting in prominent and long-lasting agonist response after washout (**Fig. 4A**). In support of this model, reintroduction of a bulky residue in the DI-DIV lateral portal (as in the case of W311F and W311A/L1439W mutants) prevents 2-APB activation (**Fig. 3B**), likely by blocking 2-APB entry into the DI–DIV fenestration. It is worth noting that multiple 2-APB binding sites, both in the extracellular and intramembrane space, have been found on members of the TRP channel family, which share similar channel architecture as Na_V_s, Ca_V_s and NALCN (59). While these findings are not direct evidence in support of our two-site model at NALCN, the ability of 2-APB to access spatially distinct sites is not unexpected given its promiscuity and membrane permeability.

To identify the putative activation site of 2-APB, we combined computational and functional approaches. We showed that there are at least two potential sites for 2-APB in the pore region of NALCN: one at the DII-DIII interface, and another at the DIII-DIV interface (**Fig. 4C–E** and **S3B**). Alanine substitutions of hydrophobic residues including F594, F1141, L1148, F1151 and Y1431 at these interfaces selectively abolish activation effect of 2-APB (**Fig. 4E**). These residues correspond to the binding residues of verapamil and cinnarizine at Ca_V_1.1 and Ca_V_1.3, respectively (**Fig. S3B**), suggesting that these binding pockets are well conserved in NALCN, despite some differences between the S6 sequences of NALCN and Ca_V_s (**Fig. S6**). Our data thus reveal the first molecularly defined drug binding sites in NALCN. While it was necessary for us to unplug NALCN’s lateral fenestrations to elucidate the 2-APB binding sites, we believe it is possible to target this site in WT channels in future drug screening efforts. We have yet to identify the speculated extracellular inhibitory site for 2-APB, possibly because multiple inhibitory sites exist on the VSDs, and occupation of more than one site is required for the inhibition of 2-APB.

### Physiological implications

Our study has illustrated that there is potential to achieve pharmacological effects beyond the occluded fenestrations of NALCN. While we have achieved this artificially in this study by reducing the side-chain volume of key bottleneck residues, it is possible that the fenestration-plugging residues may naturally adopt, or be induced to adopt different conformations, which in turn could alter the radii and accessibility of these fenestrations. In fact, there is structural evidence supported by molecular dynamics (MD) simulations that key aromatic residues along lateral fenestrations of prokaryotic and eukaryotic Na_V_s are mobile and their distinct rotamer conformations can directly gate the fenestration openings. For instance, the highly conserved phenylalanine residue in the middle of S6-DI (position 15) of Na_V_1.7 (corresponds to W311 of NALCN; **Fig. S6**) can adopt two different poses depending on local conformational changes in the PDs (**Fig. S7A**) widening or narrowing the bottleneck radius of the DI-DIV fenestration of Na_V_1.7 (46, 60). It is conceivable that these residues in NALCN which seal the lateral fenestrations in existing structural snapshots (**Fig. S1C**) can change configuration and open individual fenestrations sporadically under specific conditions. Some possibilities include ligand binding or transitions between as yet uncharacterised functional states that would expectedly trigger local conformational changes in the PDs.

The possibility of disease mutations affecting fenestration dimensions of NALCN should also be considered when interpreting our mutational data. Out of the <50 *de novo* NALCN missense mutations reported to date, there are at least two (T513N of S5-DII and F1141V of S6-DIII) that occur at residues lining the lateral fenestrations of NALCN (**Fig. 1A**). These mutations are found in the DII-DIII fenestration of NALCN and given the close vicinity of F1141 to the key bottleneck residue M1145, we expect the Phe-to-Val substitution to enlarge the fenestration portal. However, predicted tunnels through the DII-DIII interface of either T513N or F1141V are not dissimilar to that of WT, as L591 and M1145 are still restricting access (**Fig. S7C**). Nonetheless, it is worth noting that mutations of many adjacent residues including but not limited to L312I/V, V313G, F512V, L590F, V595F and L1150V have been identified in CLIFAHDD patients. As many of these *de novo* variants have drastic impact on channel function (23), it is reasonable to expect that these mutations may alter fenestration dimensions by perturbing the conformational landscape. This is conjecturally supported by our functional data that modifications in these fenestrations, for instance at position 311, can have dramatic impact on NALCN function and pharmacology (W311A/F; **Fig. 1B** and **3B**). As the study of NALCN channelopathies is still a growing field, the pathophysiological relevance of these fenestrations will hopefully become clearer in the future when more patient data become available.

### Limitations and future directions

The first limitation of this study is that our hypotheses are guided predominantly by the currently available structural information. We have identified and mutated four key residues that appear to plug individual lateral fenestrations based on apparent closed-state structures of NALCN. As proteins are dynamic entities, these static snapshots of NALCN do not report on the dynamics of fenestration dimensions. A potential issue with this approach is that residues at positions other than W311, L588, M1145 and Y1436 may take over the role of key bottleneck residues over time under different functional states. It is therefore important for future studies to use equilibrium simulations to understand the motions of fenestration-lining residues and how their conformations regulate the radii of these fenestrations. The determination of open-state or gain/loss-of-function disease mutant structures of NALCN may provide further insights into the conformational flexibility of key bottleneck residues.

The second limitation lies in the small panel of 24 compounds screened on NALCN currents. While we identified compounds such as DPBA and fluvastatin as novel efficacious inhibitors of inward current of NALCN, their minimal effective concentrations are too high (100 µM to 1 mM) for them to be considered NALCN-specific blockers. To establish a clear structure-activity relationship, it is necessary to perform high-throughput screening to gather sufficient data. However, the constitutive activity of NALCN and its absolute requirement for three auxiliary subunits to function impose significant technical barriers. An alternative approach is to synthesise and characterise a library of structural analogues of hit compounds from our work here (2-APB, DPBA, (E/Z)-endoxifen and fluvastatin) and other studies (e.g., L-703,606) (61).

### Concluding remarks

There has been enormous progress in our understanding of NALCN channelosome structure and function in the last few years, but the scarcity of NALCN-specific pharmacological tools remains a significant challenge in the field. Delineating the mechanisms underlying this pharmacological resistance is therefore crucial to unlocking NALCN’s potential as a pharmacological and therapeutic target. In this study, we have explored the hypothesis that occluded lateral fenestrations contribute to the pharmacological resistance of NALCN and our findings suggest that key bottleneck residues in this region impede drug accessibility. Despite the identification of new NALCN blockers, the low potencies of these compounds reiterate the difficulty in targeting this leak channel. Perhaps, exploring alternative strategies focusing on auxiliary subunits (UNC79, UNC80 and FAM155A) of the channelosome in the future may help overcome the present paucity in NALCN-specific modulators.

## Acknowledgements

We thank J. Colding and H. Jurewicz for technical assistance and H. Kurata for help-ful comments on the manuscript.

## Materials and methods

### Tunnel detection using MOLE 2.5

To prepare the WT channel structure for tunnel detection, we first removed the auxiliary subunits UNC79, UNC80, FAM155A and CaM from the channelosome structure (PDB 7SX3). We then used MOLE 2.5 (https://mole.upol.cz/) to detect tunnels in the structure of WT channel with default parameters as follows: minimal bottleneck radius 1.2 Å, probe radius 3 Å, surface cover radius 10 Å and origin radius 5 Å. We filtered out irrelevant tunnels that did not run parallel to the cell membrane and did not start from the central cavity. We used PyMOL to perform *in silico* mutagenesis on NALCN, and subjected mutant structures to the same procedure to detect lateral fenestrations.

### Molecular biology

We cloned the human NALCN, UNC79, UNC80 and FAM155A complementary DNAs (cDNAs) between HindIII and XhoI sites in a modified pCDNA3.1(+) vector containing a 3′-*Xenopus* globin untranslated region and a polyadenylation signal. These constructs were generated using custom gene synthesis with codon optimization for *Homo sapiens* (GeneArt, Thermo Fisher Scientific). We generated all NALCN mutants using custom-designed primers (Eurofins Genomics or Merck) and PfuUltra II Fusion HS DNA Polymerase (Agilent Technologies) or the Q5 Site-Directed Mutagenesis Kit (New England Biolabs). We verified the sequences of plasmid DNAs purified from transformed *E. coli* by Sanger DNA sequencing (Eurofins Genomics). For expression in *Xenopus laevis* oocytes, we linearised plasmid DNAs with XbaI restriction enzyme, from which capped mRNAs were synthesised using the T7 mMessage mMachine Kit (Ambion).

### Two-electrode voltage clamp

To surgically remove the ovarian lobes, adult female *X. laevis* were anaesthetized with 0.3% tricaine (under license 2014–15-0201–00031, approved by the Danish Veterinary and Food Administration). Frogs were housed and cared for by an animal facility approved by the University of Copenhagen. We then separated the ovarian lobes into smaller parts and defolliculated mechanically at 200 rpm at 37 °C. For injection, we sorted healthy-looking stage V-VI oocytes. To prepare for injection, we diluted the mRNAs of NALCN (WT or mutant), UNC79, UNC80 and FAM115A to a concentration of 1000 ng/μL and then mixed in a ratio of 1:1:1:1. Using the Nanoliter 2010 injector (World Precision Instruments), we injected 36.8 or 41.4 nL of pre-mixed RNA into each oocyte. During injection, we lined up the oocytes in OR2 medium (82.5 mM NaCl, 2 mM KCl, 1 mM MgCl2, 5 mM HEPES) and injected the oocytes at the equator region of the cell. For optimal expression, we then incubated the oocytes at 140 rpm at 18 °C for 5 days in antibiotic medium (96 mM NaCl, 2 mM KCl, 1 mM MgCl2, 1.8 mM CaCl2, 5 mM HEPES, 2.5 mM pyruvate, 0.5 mM theophylline, 50 μg/mL gentamycin and tetracycline). We performed two-electrode-voltage clamp recordings using the OC-725C Oocyte Clamp amplifier (Warner Instrument Corp, USA). The microelectrodes (borosilicate glass capillaries, 1.2 mm OD, 0.94 mm ID, Harvard Apparatus) were pulled using the P−1000 horizontal puller (Sutter Instruments) and filled with 3 M KCl and had a resistance between 0.2−1.1 MΩ. Due to NALCN’s sensitivity to the extracellular divalent cations Ca^2+^ and Mg^2+^, the oocyte was constantly perfused with Ca^2+^- and Mg^2+^-free buffer that is substituted with Ba^2+^ [96 mM NaCl, 2 mM KCl, 1.8 mM BaCl_2_ and 5 mM HEPES (pH 7.4) with NaOH], called Ca^2+^/Mg^2+^-free ND96. We acquired the data using the pCLAMP 10 software (Molecular Devices and a Digidata 1550 digitizer (Molecular devices), sampled at 10 kHz. We filtered electrical powerline interference with a Hum Bug 50/60 Hz Noise Eliminator (Quest Scientific).

For the pharmacological screening, we dissolved each tested compound to a stock solution in either Ca^2+^/Mg^2+^-free ND96 or DMSO, according to their solubility. We then diluted the stock solution further in Ca^2+^/Mg^2+^-free ND96. For the compounds that were only soluble in DMSO, the final DMSO concentration was kept at 1% or less. Due to NALCN’s pharmacological resistance, we tested the compounds at the highest concentrations that were possible while maintaining the DMSO limit.

Final concentrations of compounds tested: 1 mM 2-APB (Sigma-Aldrich), 300 μM carbamazepine (VWR), 300 μM CP96345 (Tocris Bioscience), 300 μM diltiazem hydrochloride (Alomone Labs), 100 μM L-703,606 oxalate salt hydrate (Sigma-Aldrich), 300 μM lacosamide (Sigma Aldrich), 300 μM lamotrigine (Combi-Blocks), 1 mM lidocaine hydrochloride monohydrate (Sigma-Aldrich), 300 μM nifedipine (Alomone Labs), 300 μM phenytoin sodium (VWR), 300 μM propafenone hydrochloride (Sigma-Aldrich), 300 μM quinidine (Sigma-Aldrich), 300 μM Z944 hydrochloride (Sigma-Aldrich), 1 mM diphenhydramine hydrochloride (Sigma-Aldrich), 1 mM diphenylborinic anhydride (DPBA; Sigma-Aldrich), 1 mM hydroxyzine dihydrochloride (Sigma-Aldrich), 1 mM cetirizine hydrochloride (Sigma-Aldrich), 100 µM lomerizine dihydrochloride (Sigma-Aldrich), 1 mM promethazine hydrochloride (Sigma-Aldrich), 1 mM citalopram hydrobromide (Sigma-Aldrich), 100 µM ICA−121431 (Sigma-Aldrich), 100 µM linopirdine (Sigma-Aldrich), 100 µM (E/Z)-endoxifen hydrochloride (Sigma-Aldrich) and 1 mM fluvastatin sodium (Sigma-Aldrich).

For the TEVC recordings, one oocyte (injected with either NALCN WT or mutant) was placed into the recording chamber with constant perfusion of Ca^2+^/Mg^2+^-free ND96. We then ran the first voltage protocol as a control and to check sufficient NALCN expression. Before testing compounds that were dissolved in a final DMSO concentration of 1%, we applied a Ca^2+^/Mg^2+^-free ND96 with 1 % DMSO control solution and ran the same voltage protocol to confirm that NALCN function was not affected by the DMSO content. Afterwards, we switched the perfusion in the recording chamber to the tested compound solution. To ensure that the oocyte was fully exposed to the compound, we perfused it with the compound solution for 30s and then ran the same voltage protocol again. To examine how long the effect of a compound lasted after it has been applied and washed-out, we switched the perfusion back to Ca^2+^/Mg^2+^-free ND96 and washed the oocyte for 2 min, followed by running the same protocol for a third time. To investigate the prolonged effect of 2-APB during wash-out, we continued constant perfusion with Ca^2+^/Mg^2+^-free ND96, running the same voltage protocol every 2 minutes.

### Data analysis

We analysed the recorded currents using the Clampfit 10.7 software. The raw traces were filtered at 800 Hz (Gaussian low-pass filter) and representative current traces for illustration underwent data reduction with a reduction factor of 5. We performed statistical analysis using GrahPad Prism (Version 8.4, GraphPad Software), the specific statistical tests used are mentioned where relevant.

### Molecular dynamics simulations

We extracted the coordinates of the transmembrane region of AAAA MT and WT (from I33 to M335, from A375 to S617, from M817 to F858, from H876 to E1560). We used the CHARMM-GUI Membrane Builder (62–64) to generate initial configurations of the channel embedded in a lipid bilayer and the corresponding input files. Each soluble protein–ligand system was solvated in a cubic water box that is 24 Å larger than the protein size in each direction. Distance-based ion placements were used with K^+^ and Cl^−^ ions at 0.15 M to neutralise each system. The non-bonded van der Waals interactions were truncated between 10 and 12□Å using a force-based switching method for the CHARMM Force Field (62–64). Particle-mesh Ewald summation was used for the long-range electrostatic interactions (65) and LINCS algorithm correction for hydrogen bond constraints (66). Each system was minimised for 5000 steps using the steepest descent method followed by 125□ps equilibration in the NVT (constant particle number, volume, and temperature) ensemble. Each system was then placed in a POPC homogeneous bilayer (1:1), solvated in water, and neutralized using K^+^ and Cl^-^ ions. These systems were minimised and underwent successive equilibration steps following the default CHARMM-GUI Membrane Builder equilibration protocol. Unbiased molecular dynamics (MD) simulation was conducted on the AAAA MT with 2-APB complex using Gromacs (version 2021.4) (67) with the primary aim to determine the optimal binding poses of 2-APB. This simulation was carried out under NPT conditions (constant particle number, pressure, and temperature) for ∼60 ns at a temperature of 303.15□K and a pressure of 1 bar.

### Molecular docking

We imported the cryo-EM structure of human NALCN-FAM155A-UNC79-UNC80 channelosome with calmodulin (CaM) bound (PDB ID: 7SX3) in VMD (Visual Molecular Dynamics, version 1.9.4a57), and removed the three non-conducting auxiliary subunits, CaM and all the bound lipids. Then, we mutated the four key bottleneck residues of each interface of NALCN (W311 from DI, L588 from DII, M1145 from DIII, and Y1436 from DIV) to alanine. Subsequently, we imported the AAAA MT and WT channels into Schrodinger Suite (Release 2020-4) and separately prepared both structures through the Protein Preparation Wizard module to remove original hydrogens, fill in missing side chains using Prime, cap termini, delete waters beyond 5.00 Å from protein atoms.

We chose the hypothetical binding site to be centred in the middle of the central cavity. Therefore, we generated a grid around the binding site both for AAAA MT and WT channels utilising the Receptor Grid Generation Panel with the default settings. The grid centre (193.987, 242.5975, 237.367) was chosen as the midpoint between residues T587 and Y1431. The outer grid dimensions were set to 35 Å × 35 Å × 35 Å, large enough to also include the unplugged lateral fenestrations in the case of the AAAA MT.

Using the default protocol of LigPrep (Schrödinger, LLC, 2023), we prepared the 2-APB and phenytoin structures for our docking studies. We used the program Glide (Schrödinger, LLC, 2023) to dock the prepared ligands against the receptor grids of AAAA MT and WT, keeping a rigid-receptor and flexible-ligand protocol. The ligands binding modes were ranked using the Glide extra precision scoring function. Based on the top-scoring binding poses of the different ligands, the list of potential 2-APB and phenytoin binding residues was extrapolated and used for site-specific mutagenesis.

---

**Figure S1.**
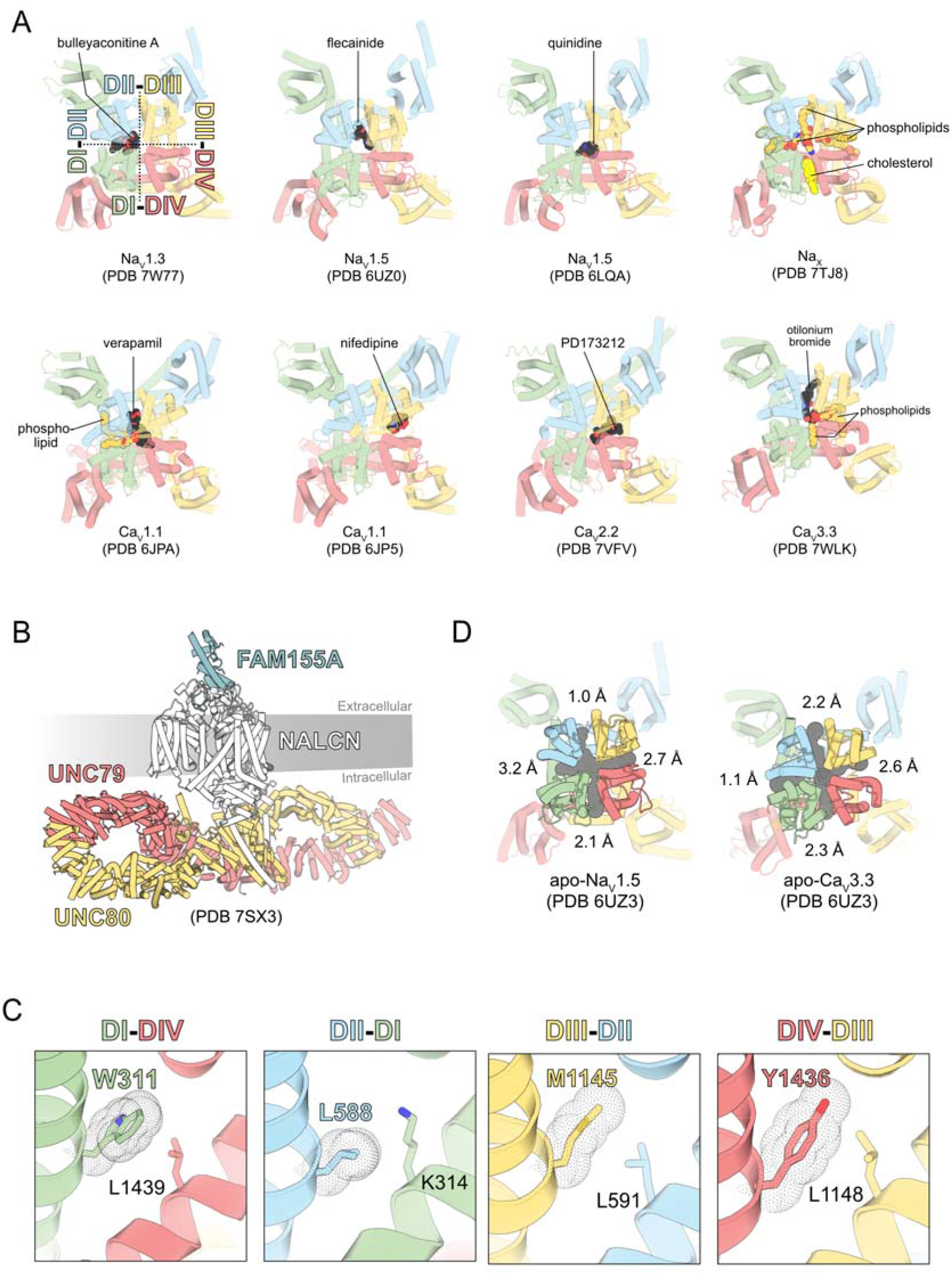
(A) Top view of Na_V_ and Ca_V_ cryo-EM structures bound to blockers and phospholipids. (B) Side view of the NALCN channelosome structure. (C) Side views of NALCN WT showing the upward configuration of key bottleneck residues occluding the lateral fenestrations of NALCN. (D) Predicted lateral fenestrations of apo-Na_V_1.5 (*top*) and apo-Ca_V_3.3 (*bottom*) channels (top view). Bottleneck radii of individual predicted tunnels are indicated accordingly.

**Figure S2.**
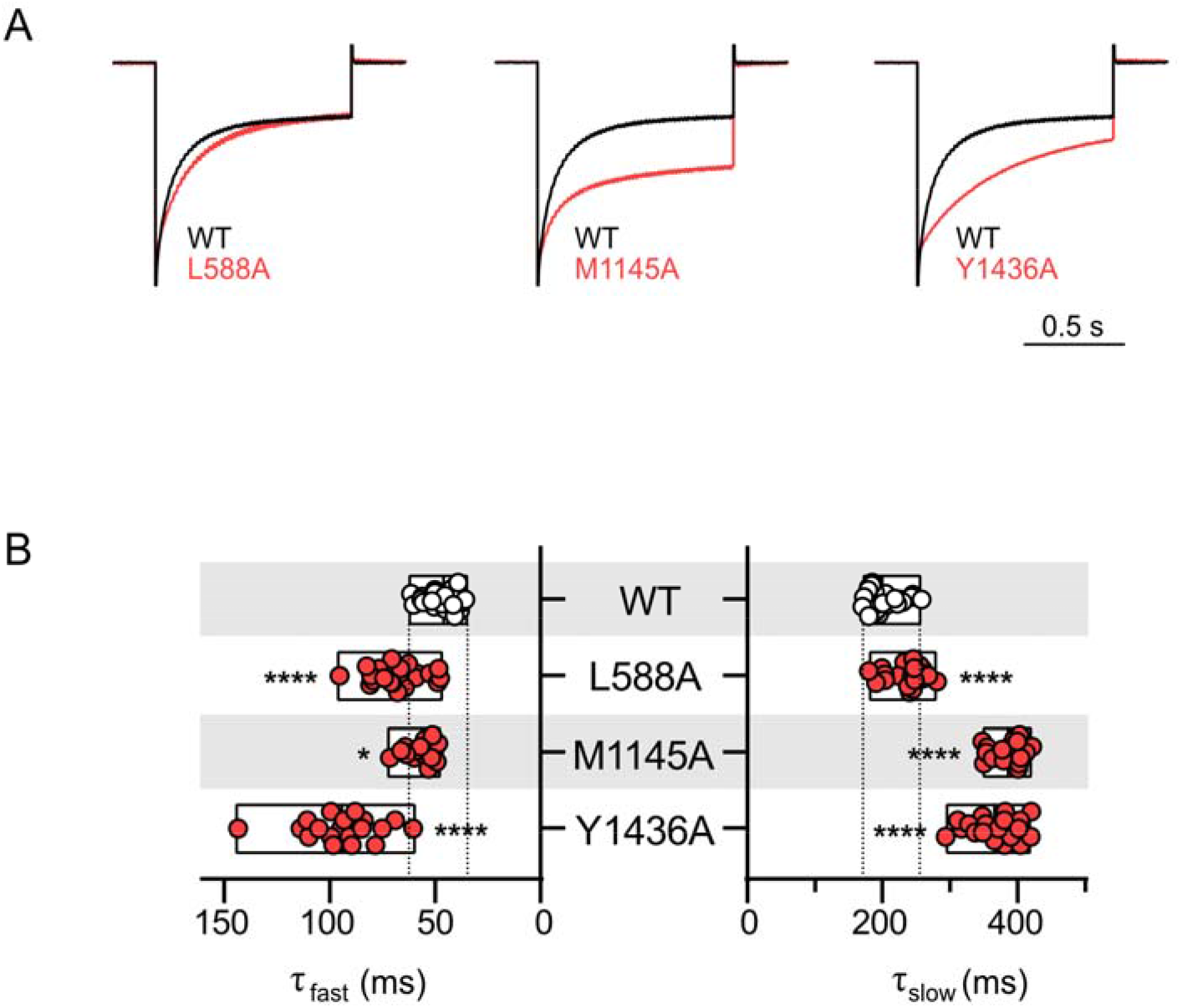
(A) Superimposed normalised current traces of WT (black) and mutants (red) at −80 mV. (B) Slow and fast time constants of hyperpolarization-elicited currents (−80 mV) for WT and mutants. \**p* < 0.05; \*\*\*\**p* < 0.0001; one-way ANOVA, Dunnett’s test (against WT). See Supplementary Table 1 for descriptive statistics.

**Figure S3.**
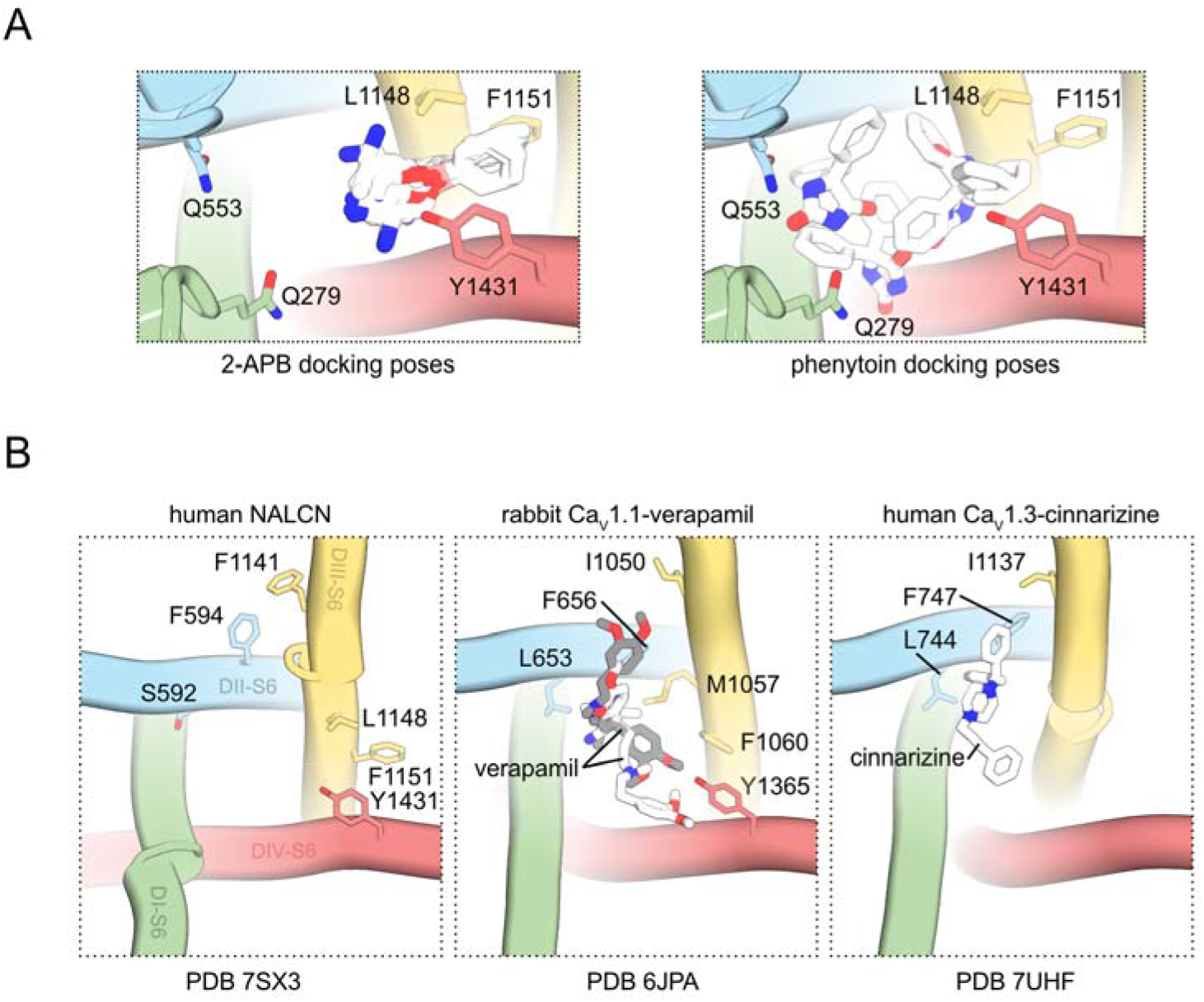
(A) *Top view*, Top-ranked docked poses of 2-APB (*left*) and phenytoin (*right*) in the pore of NALCN AAAA. (B) *Top view*, S6 segments of NALCN with residues that form the putative 2-APB binding sites within the pore region labelled (*left*). The corresponding residues in the rabbit Ca_V_1.1-verapamil (*middle*) and human Ca_V_1.3-cinnarizine (*right*) structures are highlighted.

**Figure S4.**
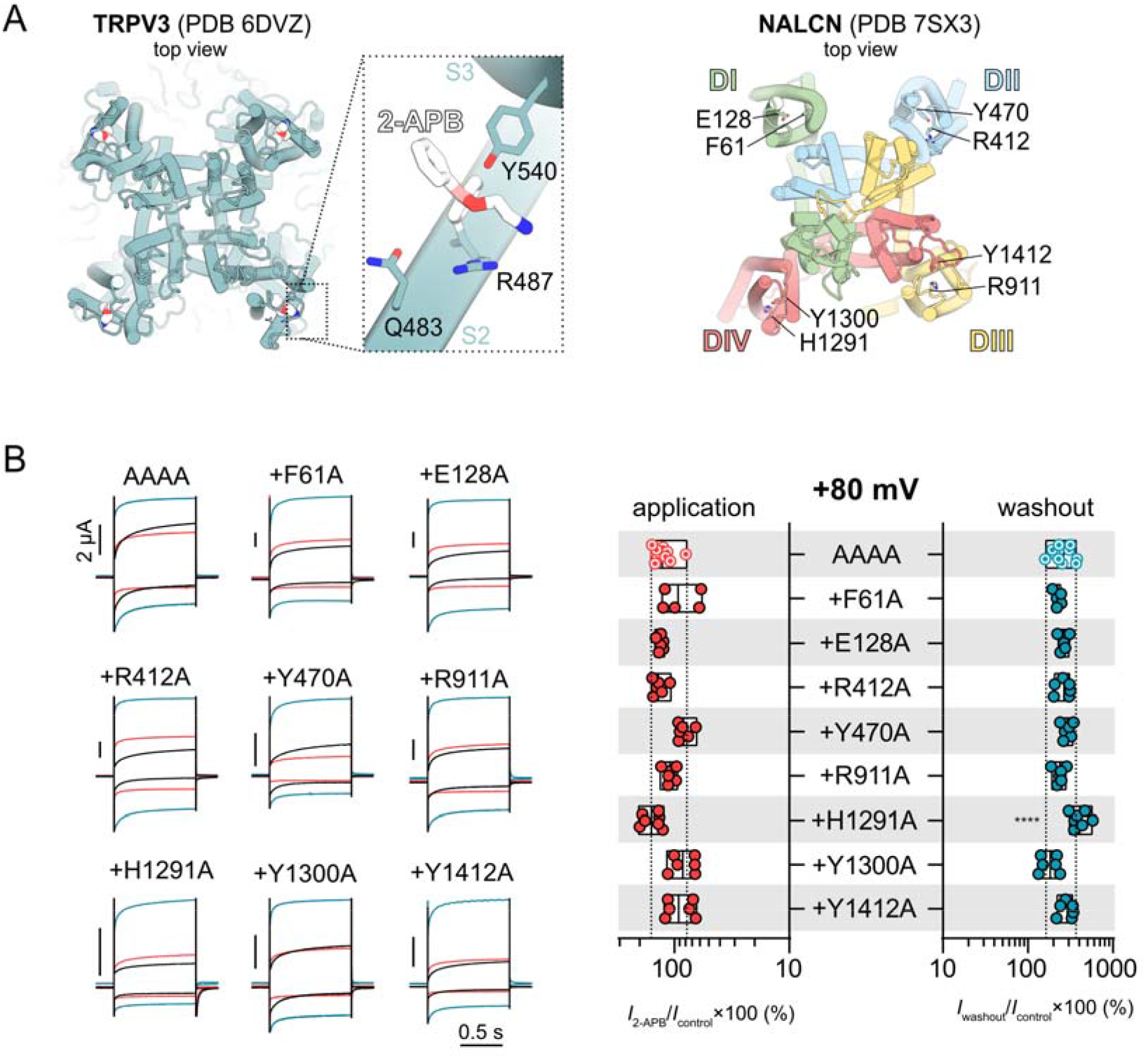
(A) *Left*, top view of the TRPV3(Y564A)-2-APB structure (PDB 6DVZ), with expanded view of 2-APB molecule in a cavity formed by the extracellular portions of S1–S4 helices. 2-APB interacts with hydrophobic and hydrophilic residues including Q483, R487 and Y540. *Right*, top view of the NALCN structure (PDB 7SX3) with hydrophobic and hydrophilic residues around the extracellular side of individual VSDs labelled. (B) *Left*, representative current traces from *Xenopus laevis* oocytes expressing various mutant channels (on the background of NALCN AAAA) in response to application of 1 mM 2-APB. *Right*, the plot shows percentage of current left for NALCN AAAA and different mutants during 2-APB application and post 2-APB washout normalised against control current elicited at +80 mV. \*\*\*\**p* < 0.0001; one-way ANOVA, Dunnett’s test (against AAAA). See Supplementary Table 1 for descriptive statistics.

**Figure S5.**
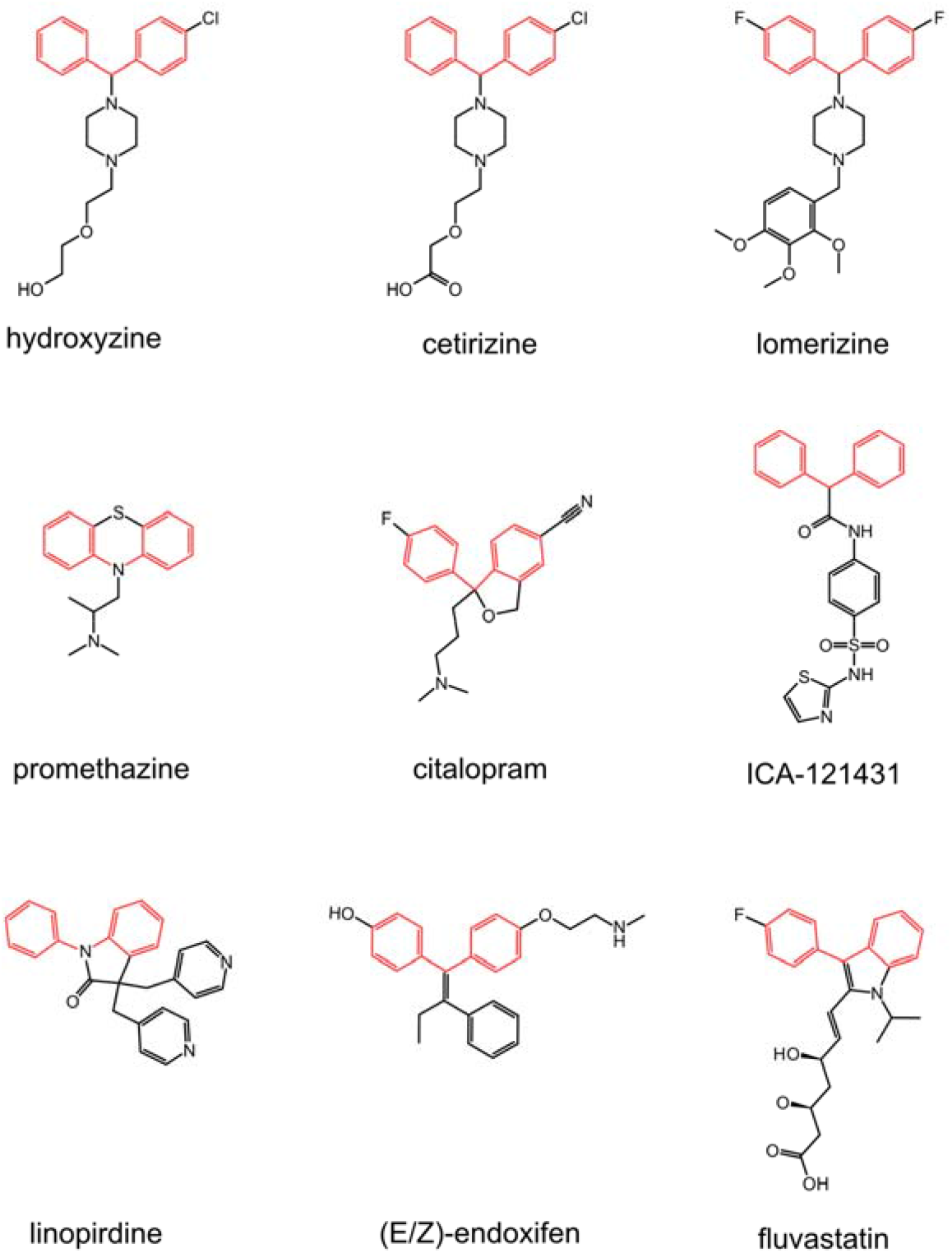
Chemical structures of compounds containing the diphenyl-methane/amine motif (highlighted in red)

**Figure S6.**
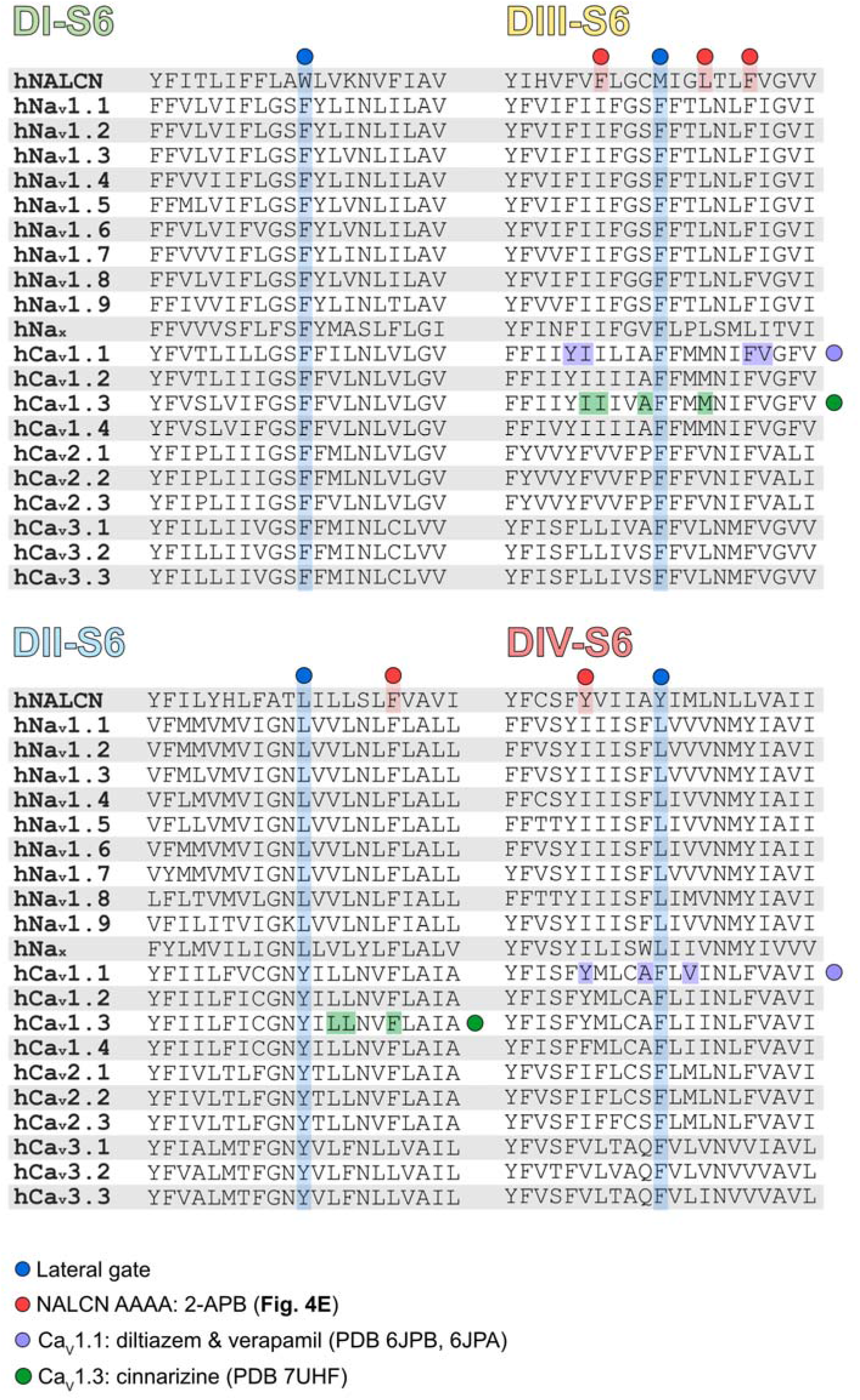
Sequence alignment of the S6 segments of human NALCN, Na_V_ and Ca_V_ channels. The residues involved in forming the lateral fenestration gates, binding and function of various compounds are shaded in different colours.

**Figure S7.**
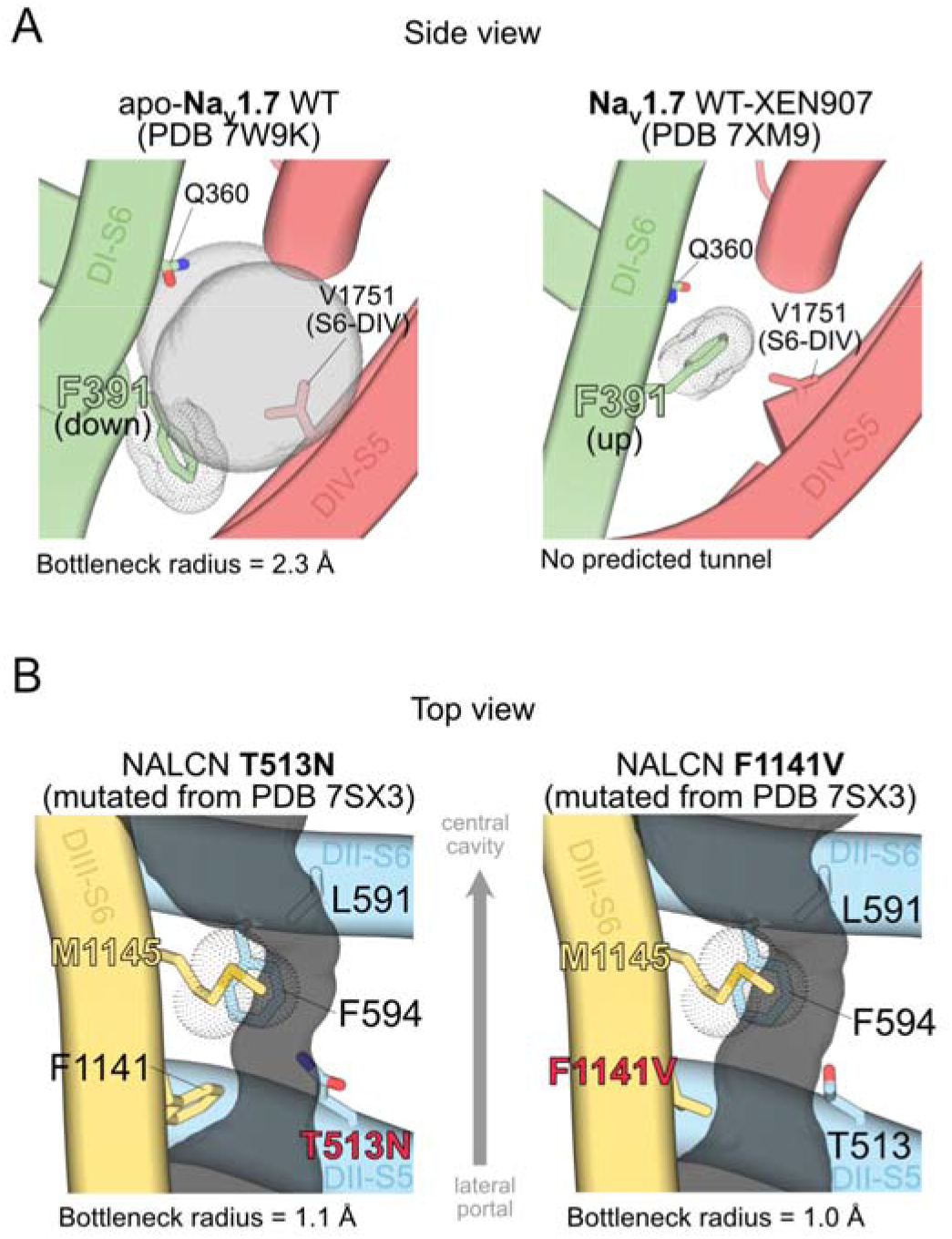
(A) Side views of Na_V_1.7 showing the flexible conformations of a highly conserved phenylalanine residue (F391) in S6-DI. F391 appears to adopt a downward configuration in the structure of Na_V_1.7 WT in the absence of bound ligands (PDB 7W9K) and an upward configuration in the presence of the Na_V_1.7 blocker XEN907 (PDB 7XM9). (B) Top views of NALCN T513N (*left*) and F1141V (*right*) channels with predicted tunnels at DII-DIII interfaces shown as grey surfaces. Key bottleneck and fenestration-lining residues are labelled. The bottleneck radius values are indicated accordingly.

